# FORGe: prioritizing variants for graph genomes

**DOI:** 10.1101/311720

**Authors:** Jacob Pritt, Nae-Chyun Chen, Ben Langmead

## Abstract

There is growing interest in using genetic variants to augment the reference genome into a “graph genome” to improve read alignment accuracy and reduce allelic bias. While adding a variant has the positive effect of removing an undesirable alignment-score penalty, it also increases both the ambiguity of the reference genome and the cost of storing and querying the genome index. We introduce methods and a software tool called FORGe for modeling these effects and prioritizing variants accordingly. We show that FORGe enables a range of advantageous and measurable trade-offs between accuracy and computational overhead.

## 1 Introduction

Assembled genomes are typically stored and understood as strings, simple sequences of base pairs. High-throughput technologies have brought an explosion of population genetics information, including from projects like HapMap [1], the 1000 Genomes Project[2] and UK10K [3]. The question is emerging: how can we use population genetics information to improve accuracy of genomic analyses? This has fueled interest in techniques that depart from a linear string as point of reference for all individuals, and toward pan-genome representations [4, 5] more inclusive of genetic variation.

While methods for including variants in the reference are growing in number [6, 7, 8, 9, 10, 11, 12], there is little or no work on how to choose which variants to include. Past studies have made such decisions in ad hoc ways, with some filtering according to allele frequency [13, 8], ethnicity [7], or both [9].

Here we examine the advantages and disadvantages of adding variants to the reference. We show that the disadvantages are important to measure, since simply adding more variation to the reference eventually reduces alignment accuracy. We suggest efficient models for scoring variants according to the effect on accuracy and “blowup” (computational overhead), and further show that these scores can be used to achieve a balance of accuracy and overhead superior to current approaches. For example, extrapolating to a whole-human DNA sequencing experiment at 40-fold average coverage, we estimate that a well-engineered augmented reference can yield about 4.8M more correctly aligned reads and 1.2M fewer incorrectly aligned compared to the linear reference. Our methods for selecting variants also reduce reference bias, a chief goals of graph genomes. Finally, we compare the accuracy yielded by our methods to that achieved using an ideal personalized graph genome. We show that our methods approach the ideal much more closely than both linear genomes – even when they are modified to contain only major alleles – and graph genomes built on different sets of variants.

These methods are implemented in a new open source software tool called FORGe. We demonstrate FORGe in conjunction with the HISAT2 [12] graph aligner and with another aligner based on the Enhanced Reference Genome [7]. But FORGe’s models and methods are suitable for any aligner that can include variants in the reference.

### Read alignment with variants

Read alignment is the process of determining each read’s point of origin with respect to a reference genome. The origin can be ambiguous and reported alignments can be incorrect [14]. Repetitive genomes and sequencing errors contribute to this problem [14, 15]. Importantly, genetic differences between donor and reference genomes also contribute. Alignments overlapping positions where the genomes differ — i.e. where the donor genome has a non-reference allele — are systematically penalized. This can (a) reduce the correct alignment’s score below the threshold considered significant by the aligner, (b) cause the aligner’s heuristics to miss the correct alignment, (c) cause the correct alignment’s score to fall below the score at a different, incorrect location. The problem is magnified in hyper-variable regions such as the Major Histocompatibility Complex (MHC) [16, 17]. It is also problematic when individuals differ dramatically e.g. if they are from distinct inbred strains [6], or when downstream analyses are vulnerable to allelic bias, such as when detecting allele-specific expression [18, 7, 19] or calling heterozygous variants [20, 21].

Augmenting the reference genome with known variants helps in two major ways. First, it reduces the genetic distance between donor and reference genomes, removing the tendency to penalize correct alignments that overlap non-reference alleles. Second, it avoids the allelic bias, also called “reference bias,” [18] that results when one donor haplotype resembles the reference more closely than the other(s).

There are many proposals for how to include and index genetic variants along with the reference genome. Two early approaches were GenomeMapper [6] and the Enhanced Reference Genome [7]. GenomeMapper came from a project to sequence many inbred strains of *Arabidopsis thaliana*, and it used a graph representation and an accompanying*k*-mer index to represent and align to a graph representing all strains. The Enhanced Reference Genome [7], which specifically addresses reference bias for allele-specific expression, included variants by taking the non-reference allele along with flanking bases and appending these “enhanced segments” to the linear reference genome. Since the resulting reference is linear, a typical read aligner like Bowtie [22] can be used.

Several studies have expanded on these ideas. deBGA [23] uses a colored De Bruijn graph [24] and accompanying hash-table index. BWBBLE [8] and gramtools [13] use an FM Index [25] with an expanded alphabet and modified backward-search algorithm to account for variants. GCSA [9] generalizes the compressed suffix array to index not a single reference but a multiple alignment of several references. HISAT2 [12] combines GCSA with the hierarchical FM Index implemented in HISAT [26]. GCSA2 [10] indexes paths in arbitrary graphs and is implemented in the VG software tool [11] which can align reads to such indexes. MuGI [27] and GraphTyper [21] use *k*-mer-based indexes.

Genome assemblies are also evolving along these lines. The GRCh37 and GRCh38 human assemblies [28, 29] include “alt loci,” alternate assemblies of hypervariable regions including MHC. Other studies suggest modifying the linear genome by replacing each non-major allele with its major alternative [30, 31]. This leverages population-level information while keeping a linear representation.

While including variants in the reference incurs a computational cost, the nature and magnitude of the cost depends on the method. For some methods, the most direct manifestation is in the size of the reference genome and index. The Enhanced Reference Genome [7] and its index both grow with the number and length of the “enhanced segments” added to cover all windows containing non-reference alleles. The number of segments required to cover a set of *k* variants within the span of a single read is 2^*k*^ *−* 1. Even if this phenomenon is isolated in a few areas of the genome (e.g. hypervariable regions), the appearance of *k* in the exponent means it can be very significant. The GCSA method[9] used in HISAT2 [12] incurs similar blowup in its “path doubling” step. We elaborate in Supplementary Note 1 and Supplementary Figures 1 and 2, including specific discussions of how the cost manifests in different methods, how it leads to more ambiguity in the reference genome which can ultimately lead to reduced accuracy, and how it can be controlled.

### Variant selection and evaluation

Past efforts that evaluated graph aligners have been selective about what variants to include in the graph, but without a clear rationale. Some included all variants from a defined subset of strains or haplotypes [6, 23, 27] or from a database such as the 1000 Genomes Project callset [2] or dbSNP [32]. In some cases, variants were filtered according to ethnicity, e.g. keeping just the Finnish 1000 Genomes individuals [9] or the Yoruban HapMap [1] individuals [7]. The ERG study (concerned with allele-specific expression) excluded variants outside annotated genes. The gramtools study [13] used 1000 Genomes variants but excluded those with observed allele frequency less than 5%. GraphTyper [21] used dbSNP variants in one experiment, excluding single-nucleotide variants (SNVs) with under 1% frequency in all populations. HISAT2’s software for selecting variants to include filters out SNVs with an allele frequency of under 10% in some cases [12].

Here we explicitly model the variants according to their effects on alignment, and we provide methods for choosing an optimal set based on those models. We apply these methods in combination with two different augmented-reference alignment methods, and compare to a range of relevant competing methods, including a linear reference with reference alleles, a linear reference with all-major alleles, and an ideal “personalized” reference that customized to fit the donor individual’s alleles (including at heterozygous positions) as closely as possible. This experimental design allows us to make statements about how our methods affect accuracy, how those effects vary with genomic region, how close the methods come to achieving ideal accuracy, and how practical current graph alignment methods are overall.

## 2 Results

### Strategy

FORGe works in cooperation with a variant-aware read aligner (*graph aligner*) such as HISAT2 [12]. Consider the alignment process as being divided into *offline* (index building) and *online* (alignment) stages. FORGe operates in the offline stage. Specifically, FORGe takes a reference genome (FASTA format) and catalog of variants and their frequencies in the population (Variant Call Format). FORGe can also use phasing information when provided in the VCF. FORGe then uses a mathematical model to score each variant according to its expected positive and negative impacts on alignment accuracy and computational overhead. The model could consider factors such as the variant’s frequency in a population, its proximity to other variants, and how its inclusion affects the repetitiveness of the graph genome. Using these scores — together with a parameter for the overall percentage or number of variants to include — FORGe outputs the top-scoring subset of variants, which can then fed to the index-building component of a graph alignment tool like HISAT2’s `hisat2-build` program. In the online stage, the aligner uses this FORGe-customized index to align the sequencing reads.

### Simulation

We used Mason 0.1.2 [33] to simulate reads (details in Supplementary Note 2). Mason simulates sequencing errors and base quality values. Mason also annotates each read with information about its true point of origin. We disabled Mason’s facility for adding genetic variants, since we simulate from already-individualized references. We classify an alignment as correct if its aligned position in the reference is within 10 nt of the true point of origin. If the aligner reports several alignments for a read, we consider only the primary alignment — of which there is exactly one per aligned read, usually with alignment score equal to or greater than all the others — when determining correctness.

### Alignment

We tested FORGe with two read alignment strategies capable of including variants in the reference: HISAT2 [12] and the Enhanced Reference Genome (ERG) [7]. HISAT2 is a practical graph aligner that we hypothesized would benefit from careful selection of genetic variants to include. The ERG is simple and compatible with linear aligners like Bowtie. We use ERG only with short unpaired reads (25 nt) to test the hypothesis that the seed-finding step of an aligner can benefit from including FORGe-selected variants. While HISAT2 can be used with unpaired and paired-end reads, we test it only with unpaired reads here. Adapting the ERG approach to paired-end alignment is probably not practical (see Discussion).

In its offline stage, HISAT2 takes a linear reference genome and a VCF file with single-nucleotide variants and indels. HISAT2 uses GCSA indexing [9] to build a graph-genome index. The resulting graph is the generating graph for all combinations of reference (REF) and included alternate (ALT) alleles. HISAT2 also provides software that, starting from a VCF file (or the UCSC “Common SNPs” track, derived from dbSNP [32]), selects a subset of variants to include. It filters in two ways. First, it excludes variants with allele frequency under 10%. Second, where variants are densely packed, it imposes artificial haplotype constraints to avoid the exponential blowup that results from considering all combinations of REF and ALT alleles. We call this the *HISAT2 auto* method.

We also tested FORGe with our implementation of the ERG [7]. ERG’s offline phase starts with a linear reference genome and a variant file. It builds an augmented reference genome by adding *enhanced segments*: reference substrings that include ALTS and flanking context. The amount of context depends on a user-specified window size, *r*, which typically equals the maximum read length. When *n* variants co-occur in a window, 2^*n*^ − 1 enhanced segments are added to cover all combinations of ALT and REF alleles. The original ERG study limited growth by considering only the leftmost *k* variants per length-*r* window, with *k* = 5 in practice. We use a variation on this limit: if a window contains more than *k* variants, we consider (a) the leftmost variant, and (b) the *k −*1 other variants with highest allele frequency according to the input VCF. Including the leftmost guarantees that each variant has its ALT included in at least one of the overlapping enhanced segments. We also set the limit higher (*k* = 15) by default. While *k* is configurable, we used the default in all experiments here. After adding enhanced segments to the reference, we indexed it with Bowtie [22]. In the online stage, we used Bowtie to align to the enhanced reference. Details on our ERG implementation are in Supplementary Note 3.

In all experiments, we ran HISAT2 with the `-k 10`, `–no-spliced-alignment`, and `–no-temp-splicesite` options. In the ERG experiments we ran Bowtie with the `-v 1` option to allow alignments with up to 1 mismatch. Note that HISAT2 is able to find alignments with mismatches, insertions or deletions, whereas Bowtie can only find alignments with mismatches. In all cases, we used Python’s `rusage` module to measure peak resident memory usage and we used the Linux `time` utility to measure real (“wall clock”) running times. All tools were run using a single thread.

### Variant models

As detailed in Methods, FORGe has two main models for ranking and selecting variants to include in the reference. First is *Population Coverage* (*Pop Cov*), which scores variants according to allele frequency. Second is *Hybrid*, which weighs both a variant’s allele frequency and the degree to which its addition would make the reference more repetitive. Additionally, we evaluated versions of these two models enhanced with a *blowup avoidance* strategy that, at variant adding time, dynamically down-weights candidates that are close to already-added variants. These versions are called *Pop Cov+* and *Hybrid+*. All of these strategies are detailed in the Methods section.

### 2.1 Chromosome 9 simulation

We tested FORGe in a series of simulation experiments. We used human chromosome 9 from the GRCh37 assembly [28]. GRCh37 was chosen to match the coordinates for the official 1000 Genomes Project Phase-3 variants [2]. We simulated sequencing reads from chromosome 9 of NA12878, a female from the CEPH (Utah residents with Northern and Western European ancestry) group studied in the 1000 Genomes Project. Specifically, we generated 10 million unpaired Illumina-like reads from each haplotype of NA12878 for a total of 20 million reads. Each read comes from one of the two haplotypes. We created a VCF file containing all single-nucleotide variants (SNVs) appearing in chromosome 9 in at least one 1000-Genomes individual, excluding NA12878 and family members. The resulting file contained 3.4 million SNVs. Details on how this set of SNVs was obtained are presented in Supplementary Note 4. We used the *Pop Cov, Hybrid, Pop Cov+* and *Hybrid+* models to score the 3.4M SNVs. The *Hybrid* and *Hybrid+* models used phasing information, whereas the *Pop Cov* and *Pop Cov+* models did not (explained in Methods). We compiled subsets of SNVs consisting of the top-scoring 0%, 2%, 4%, 6%, 8%, 10%, 15%, and 20% up to 100% in 10 point increments.

#### HISAT2

Figure 1 shows alignment rate and accuracy when using HISAT2 to align our simulated 100 nt reads to the genome indexes created with `hisat2-build`. The leftmost point (or in the case of 1c, the point labeled 0%) corresponds to a HISAT2 index with no SNVs added, i.e. a linear reference genome. The diamond labeled *Major Allele Ref* corresponds to a linear reference with all major alleles; i.e. with every SNV set to the allele that was most most frequent among CEU individuals in the filtered callset. The diamond labeled *HISAT2 auto* corresponds to the pruned set obtained by running HISAT2’s scripts. The diamond labeled *Personalized* shows results when aligning to a personalized NA12878 genome with all non-reference homozygous (HOM) alleles replaced by their ALT versions and all heterozygous (HET) SNVs added as variants, so that neither REF nor ALT are penalized at alignment time. This is not a realistic scenario, but helpful for assessing how close the tested methods come to the personalized-genome ideal. Plotted lines show results obtained when adding progressively larger subsets of SNVs to the graph genome, prioritized by model score.

**Figure 1:**
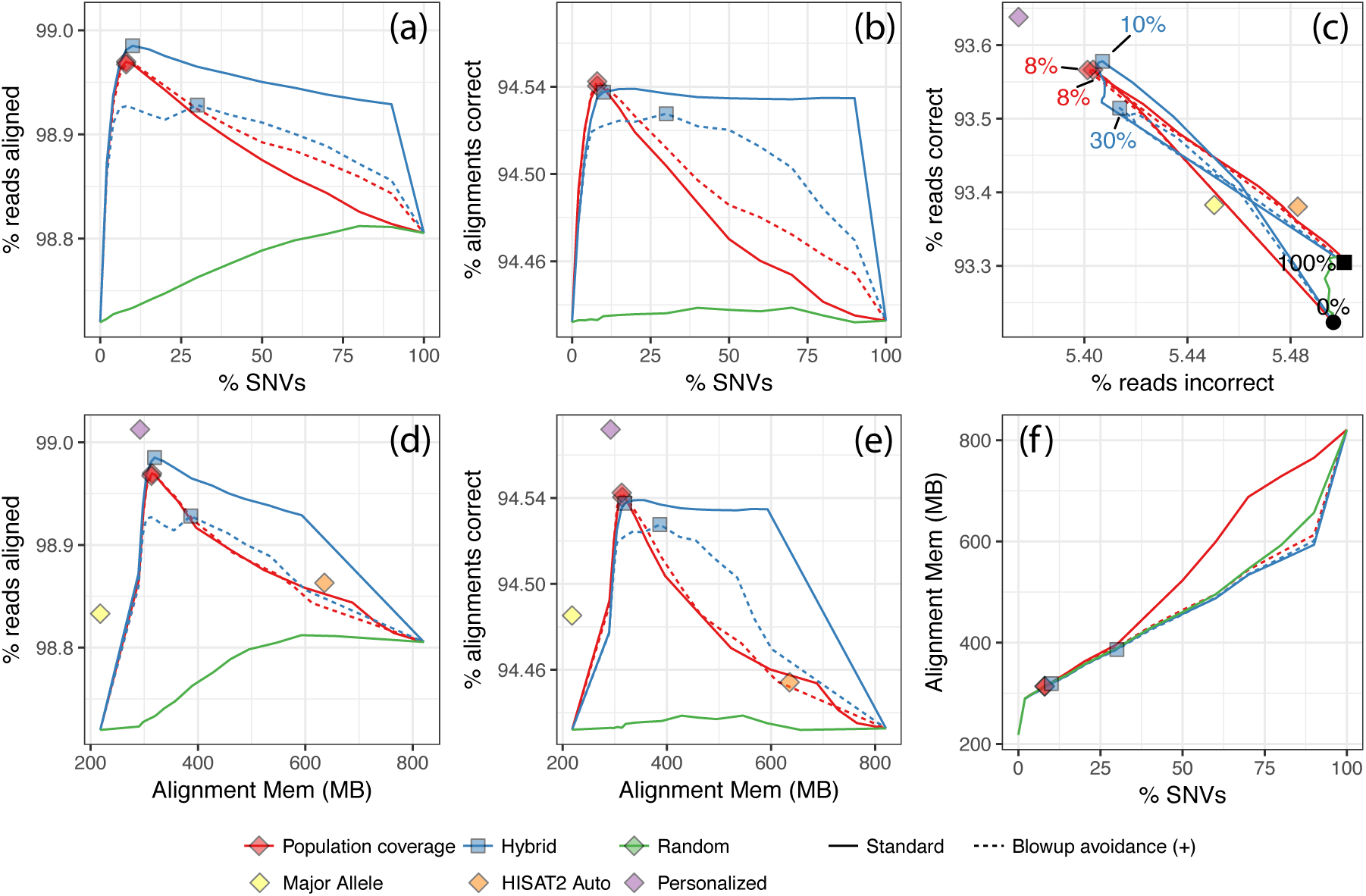
Results from NA12878 simulation for GRCh37 Chromosome 9. 100 nt unpaired reads were simulated from Chromosome 9 with NA12878’s variants included. FORGe and HISAT2 created and indexed augmented reference genomes with various variant sets. Besides the *Pop Cov* and *Hybrid* rankings, we also included a strategy that gave variants random ranks (“Random”). (a) and (d) show the fraction of reads aligned. (b) and (e) show the fraction that aligned correctly to the simulated point of origin. (c) plots a para-metric curve of the fraction of reads with a correct alignment (vertical) versus the fraction with an incorrect alignment (horizontal). Lines follow measurements made over a range of fractions of SNVs, with points for 0%, 2%, 4%, 6%, 8%, 10%, 15%, and 20% up to 100% in 10 point increments. The diamond labeled *HISAT2 auto* is an augmented genome produced using HISAT2’s pruning scripts. The diamond labeled *Major allele ref* is a linear reference with all positions set to the most frequent allele. Other diamonds indicate the SNV fraction maximizing *y* − *x*, where *y* is the fraction of reads aligned correctly and *x* is the fraction aligned incorrectly. The *HISAT2* and *Major allele* diamonds are excluded from panels (a), (b) and (f) because there is no clear way to measure the fraction of variants included by these methods. The black filled circle and square in panel (c) represent measurements when 0% and 100% of variants are included, respectively.

Figures 1a and 1b show alignment rate and fraction of alignments that are correct (henceforth “correctness”) as a function of the number of SNVs included in the genome. For all models except *Hybrid+*, peak alignment rate and correctness occur in the 8–12% range of SNVs included. All the FORGe models at their peak achieve higher alignment rate and correctness than the major-allele and HISAT2 methods. When greater fractions of variants are included — more than around 12% — alignment rate and correctness generally decrease. Correctness eventually decreases to a level only somewhat higher than that achieved by the linear reference, showing that alignment suffers when too many variants are included. Figures 1d and 1e are similar to 1a and 1b but show alignment rate and correctness as a function of HISAT2’s memory footprint at alignment time. While FORGe’s models at their peak have a roughly 50% larger memory footprint than the linear references (both major-allele and reference-allele), they use roughly half the memory of the “HISAT2 auto” method.

Figure 1c plots a point or a parametric curve for each indexing strategy and model. The vertical axis is the fraction of reads (not alignments) that aligned correctly, and the horizontal axis is the fraction of reads that aligned incorrectly. Notable points on the curves are labeled with the fraction of SNVs included. Diamonds mark points on the curves with maximal *y −x*, where *y* is fraction correct and *x* is fraction incorrect. This is a combined measure for alignment rate and accuracy, and maximal values are reached in the 8–10% range of SNVs included (except *Hybrid+*, which peaked at 30%). The best-performing are superior to (above and to the left of) the linear-genome methods, the “HISAT2 auto” method, and to the genome obtained by adding all of the SNVs (labeled 100%). The best-performing graph genomes come much closer to the personalized-genome ideal than the other methods.

It is notable that the alignment rate curves in Figure 1a,b,d and e eventually trend downward. Like most read aligners, HISAT2 uses heuristics to limit the effort spent aligning reads to many repetitive regions of the same reference genome. HISAT2 is unusual in that when a read has too many repetitive alignments, it will abort and leave the read unaligned. Bowtie does not have this heuristic; rather, Bowtie chooses one best-scoring alignment to report even when the read has many repetitive alignments. Because of this, HISAT2’s alignment rate decreases as more variants are included and the genome becomes more repetitive.

A known drawback of graph aligners is that accuracy and overhead can suffer when many variants co-occur in a small window of the genome. To measure the impact this has on FORGe’s models, we also plotted results using *blowup avoiding* versions of the *Pop Cov* and *Hybrid* models (Figure 1, dotted lines), called *Pop Cov+* and *Hybrid+*. These versions will, when selecting variants to add, deprioritize variants that are near already-added variants. We observed that blowup avoidance had a minimal impact on the shape of the *Pop Cov* curve; e.g. Figure 1d & e shows the solid and dotted lines for *Pop Cov* on top of each other. Notably, blowup avoidance did cause the alignment memory to increase more slowly with respect to the number of added variants for the *Pop Cov* ranking (Figure 1f). For the *Hybrid* model, blowup avoidance did not change the relationship between memory footprint and number of variants added (Figure 1f) and had an adverse effect on alignment rate and correctness. This is likely because the *Hybrid* model already takes clustered variants into account in its *k*-mer counts.

We repeated these experiments for paired-end reads (Supplementary Figure 3) and the results closely followed those in Figure 1. Alignment rate and accuracy both increased when using paired-end reads, since an accurate alignment for one end can “rescue” the other in the presence of ambiguity. Peak accuracy (maximal *y* − *x*) was achieved at the same SNV fraction except in the case of the Hybrid ranking, which peaked at 15% rather than at 10%.

We also repeated these experiments for reads simulated from Yoruban (YRI) individual NA19238, also sequenced in the 1000 Genomes Project (Supplementary Figure 4). As we did for NA12878, we excluded variant calls for NA19238 and family members before providing variants to the model for scoring. These results also closely followed those in Figure 1, with accuracy and recall peaking at a somewhat higher percentage of variants included (15% for YRI compared to 8-10% for CEU), likely due to YRI’s greater divergence from the reference. We return to this in the Discussion.

Finally, we repeated the unpaired NA12878 experiment including both SNVs and indels in the FORGe analysis (Supplementary Figure 5). Whereas previous experiments modeled and scored 3.4M SNVs, here we modeled and scored 3.4M SNVs and 131k indels, composed of 49k insertions ranging in length up to 411 nt and 82k deletions up to 92 nt. Given these variant scores, we selected top-scoring fractions, built indexes, simulated reads from NA12878 (both SNVs and indels included) and performed alignments as before. When assessing correctness of the resulting read alignments, we took coordinate shifts due to indels into account. Overall the results are similar to those in Figure 1. While there is a slight drop in peak alignment and correctness rate, the rates varied over a wider range of percentages relative to the SNV-only experiment. Maximal *y −x* occurred at slightly higher variant fractions relative to the SNV-only experiment: 10% for *Pop Cov* and *Pop Cov+*, 15% for *Hybrid* and 30% for *Hybrid+*.

#### Enhanced Reference Genome

Figure 2 shows alignment rate and correctness when using Bowtie [22] to align simulated 25 nt reads to enhanced references constructed with the ERG method [7]. We used shorter reads and configured Bowtie to find alignments with up to 1 mismatch (`-v 1`) to mimic the seed alignment step of seed-and-extend aligners. Unlike HISAT2, Bowtie always reports an alignment if one is found, regardless of how repetitively the read aligns. Consequently, the alignment rate shown in Figure 2a and d strictly increases as variants are added to the graph. Apart from that, the results reinforce those from Figure 1. Peak alignment rate occurs at a relatively small fraction of SNVs (6-20%). As more variants are added, decreases eventually decreases, though the *Hybrid* ranking does not suffer this drop until over 70% of SNVs are included. The alignment-time memory footprint of the best-performing FORGe indexes is higher than that of the linear reference; e.g., including the top 6% of *Pop Cov+*-scored SNVs increases the footprint 29%, from 127.9 MB to 165.0 MB. But it is a fraction of the size of the index when 100% of variants are included (1.87 GB). Blowup avoidance (Figure 2, dotted lines) had a somewhat minor effect on alignment rate and correctness for *Pop Cov*, and a clear negative effect for *Hybrid*. On the other hand, it slowed the rate of index growth for both models at low and intermediate fractions of SNVs (Figure 2f).

**Figure 2:**
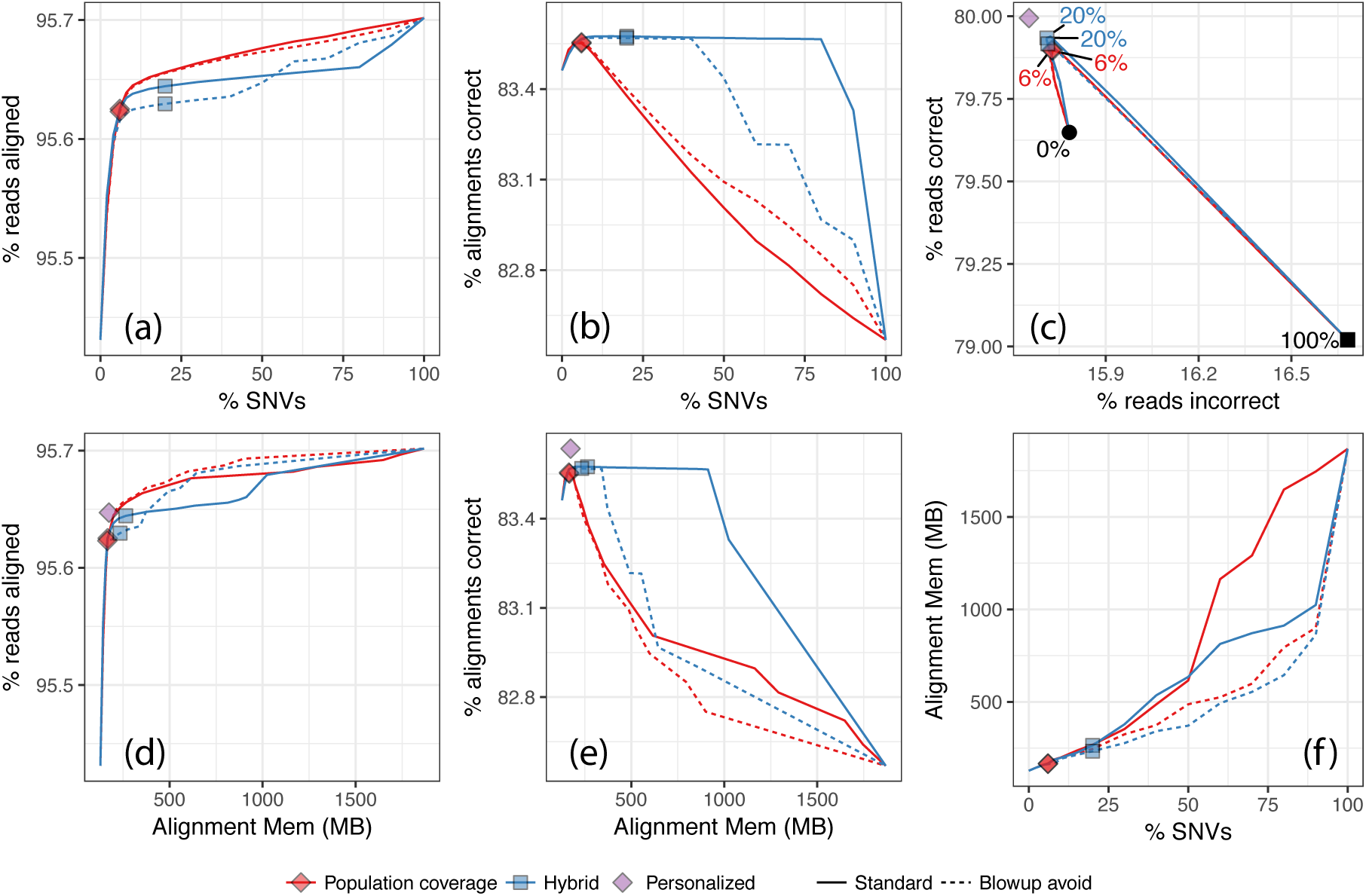
Results from aligning simulated 25 nt unpaired reads to GRCh37 chromosome 9 using the ERG+Bowtie approach. Panels and plots have the same meaning as for Figure 1 except that *HISAT2 auto* and *Major allele ref* diamonds are omitted here.

### 2.2 Stratification by variant density, variant rarity, and repetitiveness

Figure 1c showed that when we move from 0% to 8% of variants included in the augmented reference, the number of correct alignments increases by about 0.4 percentage points (as a fraction of reads) and the number of incorrect decreases by about 0.1 points. Though these may seem like small differences, in a study with 1.2 billion reads — approximately the number of unpaired 100 nt unpaired reads required to cover the human genome to 40-fold average depth — this would yield about 4.8M more correctly aligned reads and 1.2M fewer incorrectly aligned.

Still, we hypothesized that certain read subsets might be affected more dramatically by the inclusion of variants. To this end, we measured alignment rate and correctness when we varied the number of alternate alleles overlapped by a read (3a-c), whether the alternate allele was common or rare (3d-f) and what kind of genomic region or repeat the read originated from (3g-i). The measurements studied here are the same as those presented in Figure 1, but filtered as described below.

Figures 3a-c show alignment rate and correctness stratified by the number of non-reference SNVs overlapped by a read. To obtain these subsets, we first removed reads originating from reference-genome regions deemed repetitive by DangerTrack [34] (score over 250). We did this after finding that these regions had a combination of low SNV density and repetitive content that caused the 0-SNV stratum to behave very differently from the others. Reads containing 1 or more SNVs undergo a rapid increase in alignment rate and correctness from 0% to 10% of SNVs. Beyond 10%, all strata experience a slow decrease in alignment rate and correctness up to 100% of SNVs added. The 0-SNV stratum has decreasing alignment rate and correctness across the whole range, as expected since the addition of variants cannot help (since the reads lack alternate alleles) but can harm alignment by increasing the repetitiveness of the reference. Strata with more SNVs experience a more dramatic rising-and-falling pattern; for the 3-SNV stratum, alignment rate varies from about 80–98%. While curves for the various strata have different shapes, all peak at a relatively low SNV fraction: 20% or lower.

**Figure 3:**
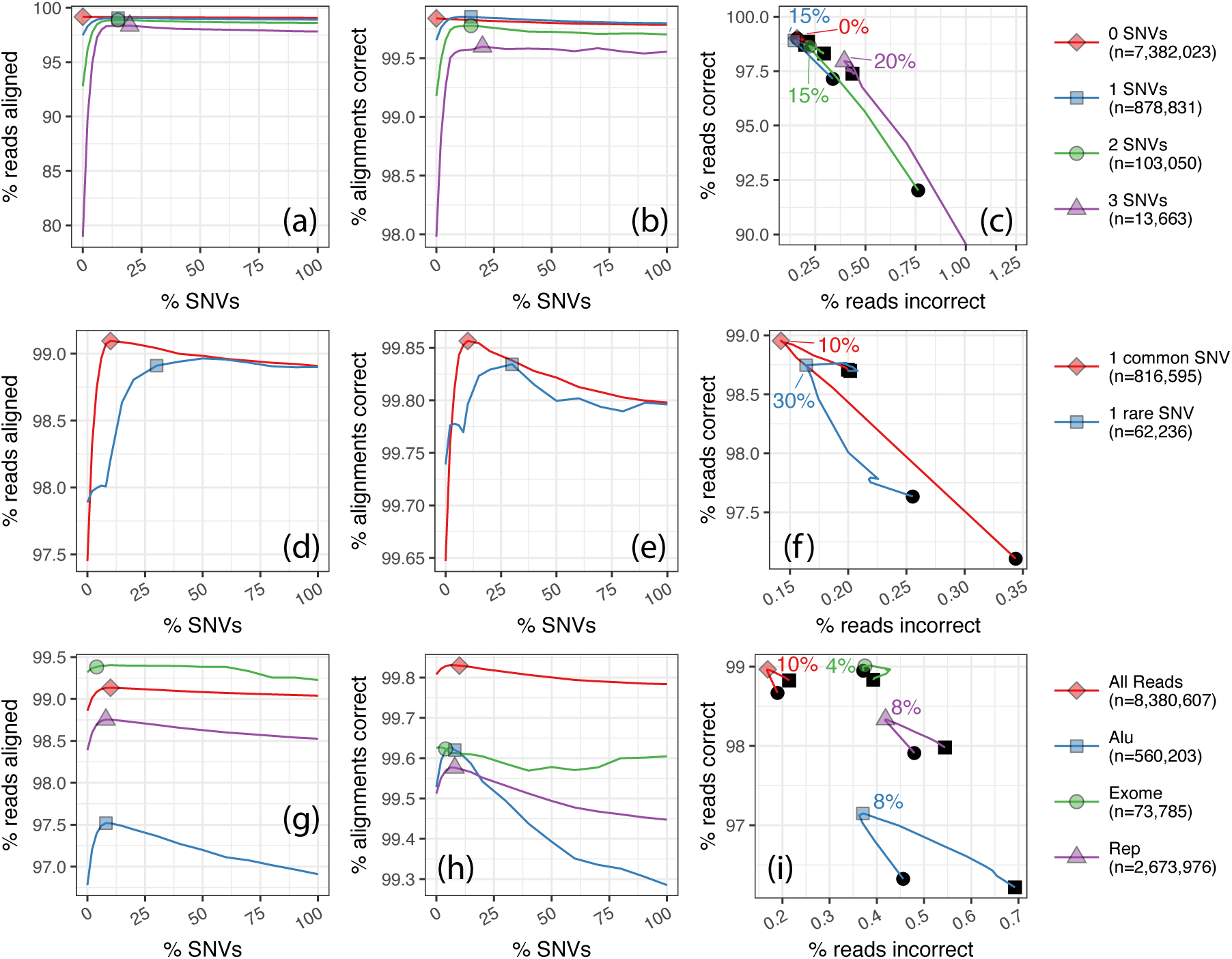
First row: Results for simulated reads stratified by the number of SNV alternate alternate alleles overlapped by the read. Reads overlapping regions with high DangerTrack [34] score — indicating the regions are difficult to align to — are omitted. Second row: Results for simulated reads overlapping exactly one common alternate allele (and no other alternate alleles) and reads overlapping exactly one rare allele. Reads overlapping regions with high DangerTrack [34] score, indicating the regions are difficult to align to. Third row: Results for simulated reads stratified by region of origin. Regions examined are: regions labeled with the “Alu” family by RepeatMasker, regions captured by the Nextera exome sequencing protocol (“Exome”), and regions labeled with any repeat family by RepeatMasker (“Rep”).

Figures 3d-f show alignment rate and correctness for reads containing a single rare SNV allele (1000 Genomes frequency < 0.5) versus reads containing a single common SNV allele (≥ 0.5). In both cases, we considered only reads with a single non-reference allele. Rare-SNV reads peak lower and at a higher SNV fraction than common-SNV reads for both alignment rate and correctness (Figures 3d-f). This is expected, since the *Pop cov* model prioritizes common over rare SNVs. In other words, by the time a rare variant is added, many common variants have already been added, making the genome more repetitive.

Figures 3h-j show alignment rate and correctness for reads stratified by feature of origin. We analyzed reads originating from (a) RepeatMasker-annotated repetitive regions(http://www.repeatmasker.org), (b) RepeatMasker-annotated “Alu” repeats, (c) regions captured by the Nextera exome sequencing protocol, and (d) all reads. Reads from repetitive regions generally had lower alignment rate and correctness compared to all reads. As before, alignment rate and correctness curves peaked at low SNV fractions: 10% or lower. Reads from more repetitive features were more sensitive to the number of variants included in the reference, as evidenced by the vertical spans of the curves.

In a related experiment, we examined the graph genome’s effect specifically on the hypervariable MHC region. We simulated reads from NA12878 Chromosome 6 and used HISAT2 to align to both a linear and a graph genome augmented with the top-scoring 10% of SNVs. We visualized the read-alignment pileup in the hypervariable MHC region using IGV [35] (Supplementary Figure 6). Qualitatively, the pileup for the augmented reference looks superior — with more coverage in variant-dense regions and with more even overall coverage — to the pileup for the linear reference.

### 2.3 Ethnicity specificity

We also studied how ethnicity-specific augmented references, advocated in other studies [36, 37, 38], can improve alignment. We used FORGe to select variants from two lists: one with variants drawn from and scored with respect to the overall 1000-Genomes phase-3 callset, and another drawn from and scored for just the CEU individuals. In both cases, variants private to NA12878 and family members were excluded and reads were simulated from NA12878.

Figure 4 shows alignment rate and correctness when aligning to CEU-specific and pan-ethnic references. As expected, the CEU-specific reference yielded higher alignment rate and correctness. CEU-specific curves also peaked at lower numbers of SNVs compared to pan-ethnic. However, the differences were only a few hundredths of a percentage point and cover only a small fraction of the remaining distance to the ideal point. Looking at this another way, if we extrapolate the results to a whole-genome DNA sequencing experiment with 40-fold average coverage, around 250,000 alignments would be affected. We return to these small differences in the Discussion section.

**Figure 4:**
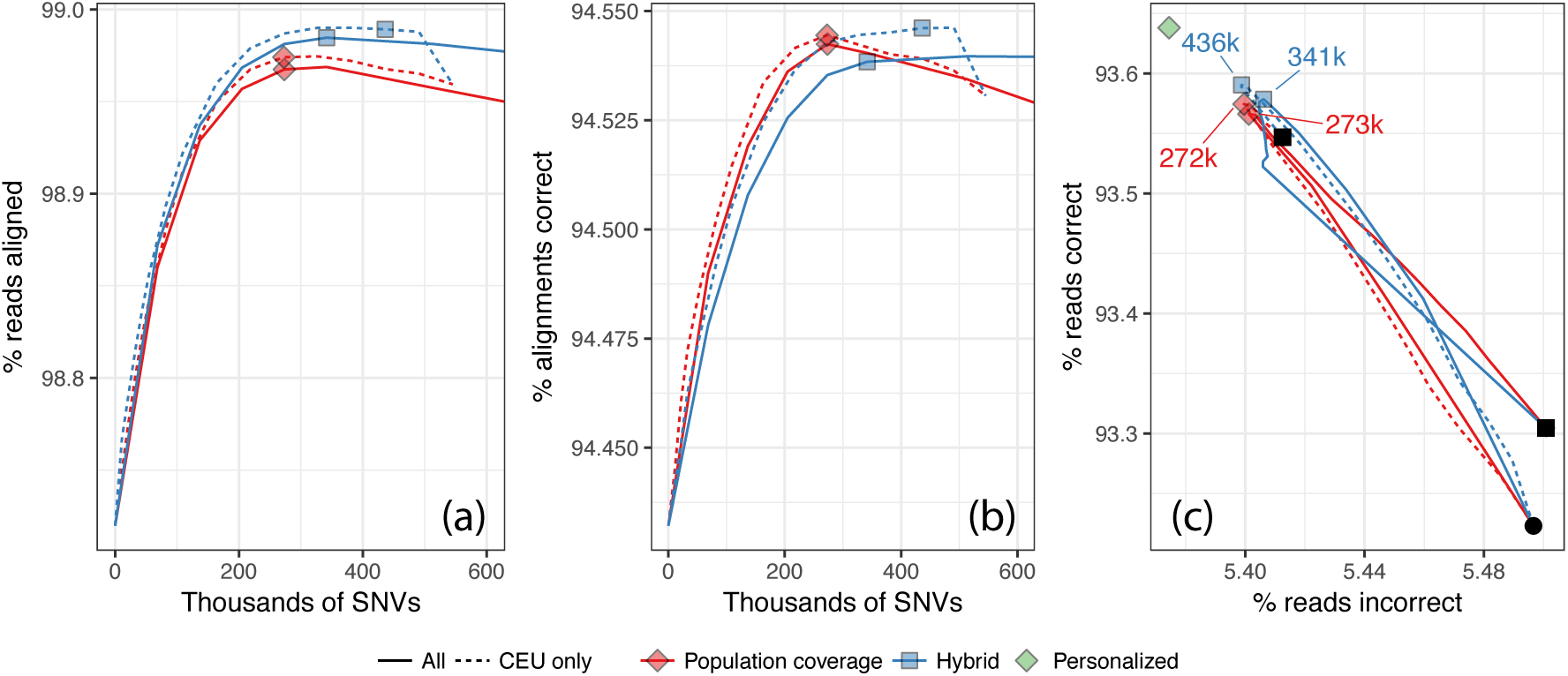
Results from chromosome-9 NA12878 simulation when using an ethnicity-specific (“CEU”) versus a pan-ethnic (“All”) augmented reference. Reads are 100 nt and unpaired and the plots have similar axes as the plots in Figure 1 panels (a)–(c). Since we are assessing two ethnicities, each with a different total number of variants, the horizontal axes in panels (a) and (b) and the peak points in panel (c) are labeled with absolute number of variants included rather than percentages.

### 2.4 Whole human genome

#### Simulated reads

To show our methods generalize to whole genomes, we repeated experiments like those presented in Figure 1 using the full GRCh37 reference. We gathered 80.0 million SNVs from the Phase-3 callset of the 1000 Genomes Project [2]. We used FORGe’s *Pop Cov+* model to score SNVs and compiled subsets consisting of the top-scoring 2%, 4%, 6%, 8%, 10%, 15%, and 20% up to 100% in 10 point increments. We built graph-genome indexes for each using HISAT2. We used the *Pop Cov+* model because the others required excessive time and/or memory; specifically, the *Pop Cov* model (without blowup avoidance) produced a set of variants that HISAT2 was unable to index in a practical time and space budget (Supplementary Note 5) and the *Hybrid* and *Hybrid+* models required excessive time for the step that generates the FASTA file for *G* due to exponential blowup (Supplementary Note 6).

Figures 5a & b plot HISAT2 alignment rate and correctness as a function of the SNV fraction. We aligned 20 million 100 nt unpaired reads from simulated from NA12878. We omitted NA12878 and family members from variant selection. Results using the ideal personalized index are also shown for comparison. Maximal *y* − *x*, where *y* is the fraction of reads aligned correctly and *x* is the fraction aligned incorrectly, occurred at 10% of SNVs (Figure 5c). Interestingly, the maximal point does not approach the personalized-genome ideal point as closely here as it did for the chromosome-9 experiment (Figure 1). This seems to be due to the added ambiguity that comes when variants in all non-chromosome-9 portions of the genome are added (Supplementary Figure ad7).

**Figure 5:**
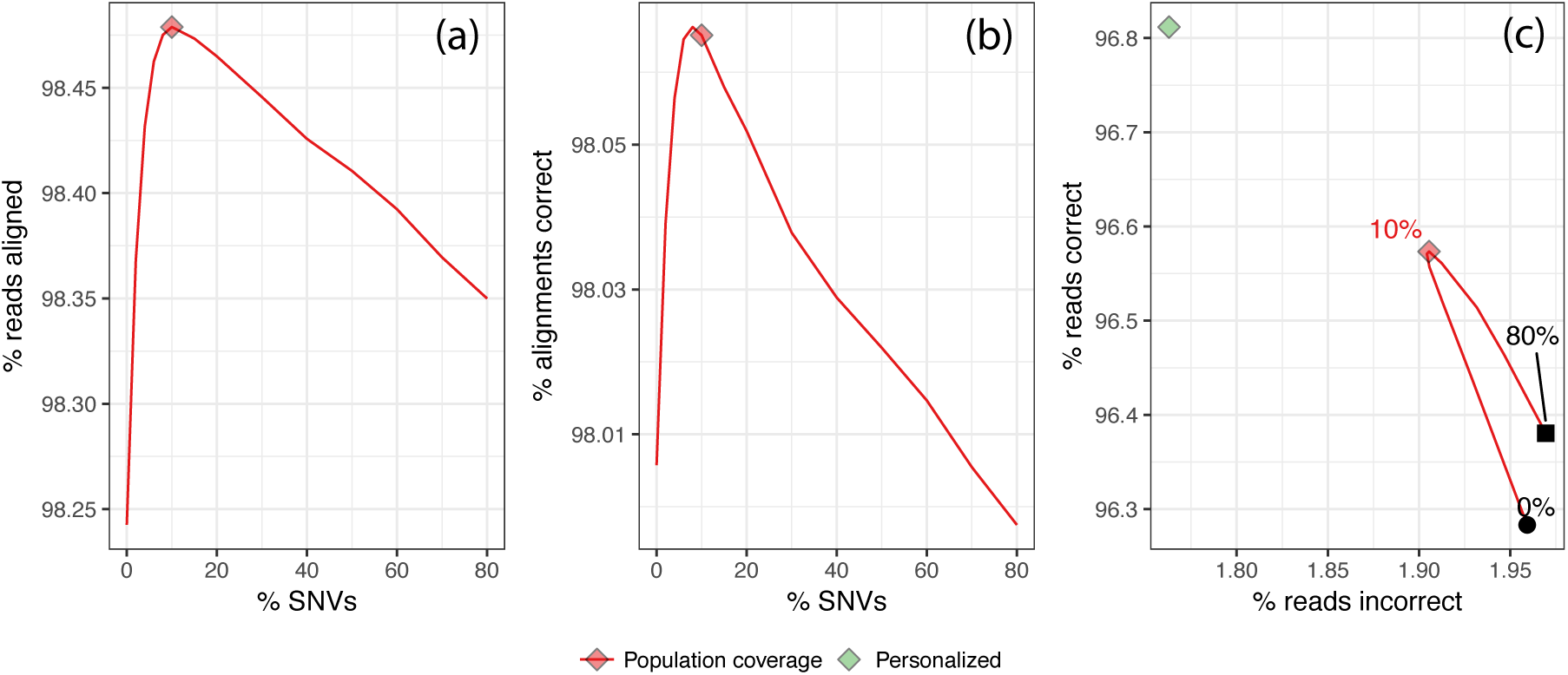
Results from aligning NA12878-simulated reads to HISAT2 graph genome for the whole GRCh37 genome. Variants were selected using FORGe’s *Pop Cov+* model. Plots have the same axes as the plots in Figure 1 panels (a)–(c). The green diamond in panel (c) shows the result when aligning to a personalized graph genome with exactly the individ-ual’s variants.

#### Platinum reads, SNVs

We conducted further experiments using a set of 1.57 billion real 100 nt unpaired sequencing reads from the Platinum Genomes Project [39] (accession: ERR194147). Like the simulated reads, these also come from NA12878. We gathered a set of 80.0 million SNVs from the 1000 Genomes phase-3 callset, omitting variants private to NA12878 and family members. We again used the *Pop Cov+* model to select variants.

We cannot assess correctness since the reads were not simulated. Following a prior study [40], we measured the number of reads that align uniquely — where HISAT2 reported exactly one alignment — versus the number that aligned perfectly, matching the reference exactly with no differences. The goal was to capture the variant-inclusion trade-off; we hypothesized that adding more variants will remove the alignment-score penalty associated with known genetic variants (increasing the number of perfect matches) without increasing reference ambiguity (decreasing the number of unique alignments). As shown in Figure 6a, the points that achieved the peak number of unique plus perfect alignments corresponded to 30% of the SNVs. This fraction is higher than most of our simulated results, perhaps due to the fact that unique-plus-perfect is an imperfect proxy for correct-minus-incorrect (Supplementary Figure 8).

**Figure 6:**
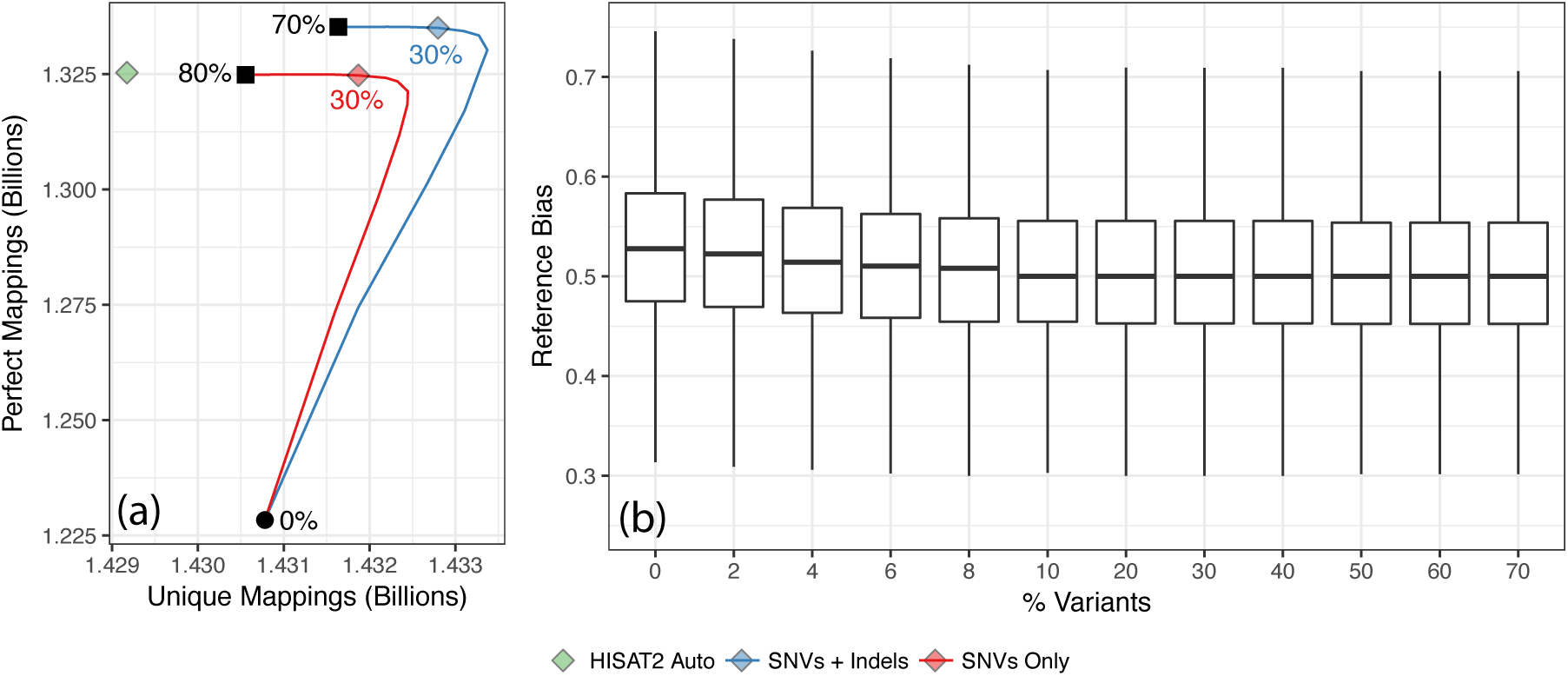
(a) Perfect/unique alignment results when aligning real reads. The blue curve is parametric, as a function of the fraction of variants included from 0% (bottom left) to 80% (top). The green diamond marks the number perfect and unique mappings for HISAT2’s custom variant pruning script applied to the set of SNVs and Indels. Graph genomes were built for SNVs alone (red) and for SNVs and Indels (blue), both ranked with the *Pop Cov+* strategy. Blue and red diamonds mark the fractions that achieved the highest sum of unique and perfect alignments. (b) Allelic bias for the 2.07 M heterozygous SNVs that met a minimum coverage threshold of 25 in all experiments. Whiskers show the 5th-to-95th percentile range.

#### Platinum reads, SNVs and indels

To highlight the effect of including indels in the reference, we repeated the previous experiment but using both SNVs and indels from the 1000 Genomes phase-3 callset. Specifically, we gathered 83.1 million variants, both SNVs and indels, but omitting variants private to NA12878 and family members. We again used the *Pop Cov+* model to select variants. We again plotted the number of reads that aligned uniquely versus the number that aligned perfectly (Figure 6a). The graph genome built from both SNVs and indels achieved peak unique+perfect at 30% of variants, like the graph built from SNVs alone. However, at every percentage it yields more unique and perfect alignments.

#### Reference bias

We measured how reference bias varies with the fraction of variants included. We analyzed the alignments of the ERR194147 reads to the whole human genome with both SNVs and indels included in in reference. Figure 6b shows a series of boxplots summarizing bias at a set of 2.07 million HET SNVs called in NA12878 by the Platinum Genomes Project [39]. The set of 2.07M HETs was chosen by taking all HETs covered by at least 25 reads in all of our experiments. Each boxplot summarizes the fraction of REF alleles (*REF/*(*REF* + *ALT*)) at the HET site for all 2.07M HETs. As expected, bias decreased as more variants were included. The decrease plateaued at 10–20% of variants. Beyond 20%, including more variants did further reduce bias, but only slightly. From 20% to 70% of variants the mean decreased by only 0.00011. This is consistent with previous results showing that most of the benefit is achieved at a small fraction of variants.

#### HLA typing accuracy

Finally, to measure FORGe’s effect on downstream results, we measured how HLA typing recall vary with the fraction of variants included in the reference. We used the same NA12878/ERR194147 alignments evaluated in previous sections, extracted alignments in the MHC region, then provided those alignments to the Kourami [41] HLA typing tool to make HLA calls. We repeated this with indexes for all the same variant-inclusion fractions evaluated previously. More details on the HLA typing methodology are described in Supplementary Note 7. In comparison with linear genome, HLA typing recall and accuracy increased substantially when the highest-scoring 10% of SNVs were included in the augmented reference. Recall and average coverage plateaued at larger SNV fractions (Supplementary Figure 9). Overall, we see that — as we observed in other results — HLA allele recall benefits from the addition of a carefully-chosen fraction of variants, and that a fraction of only 10% is sufficient to achieve peak recall.

## 3 Methods

FORGe works in cooperation with a variant-aware read aligner such as HISAT2 [12] or the ERG [7]. The strategy has two stages. In the *offline* stage, FORGe selects variants to include in the augmented reference based on a variant model — which predicts the pros and cons of including a variant — and a variant limit. The model and limit together constitute a *variant inclusion strategy* (VIS) that aims for a balance between accuracy and overhead. Once variants have been selected, the aligner software is used to create an index of the augmented reference. The second stage is an *online* stage where the read aligner aligns reads to the augmented reference using the index.

### 3.1 Offline stage

Inputs to the offline stage consist of (a) a reference genome, (b) variants in VCF format, (c) a VIS, and (d) a window size *s*. The variant inclusion strategy (VIS) consists of a variant model and a limit on the number or fraction of variants to include. The VIS is the user’s most direct means for balancing blowup and alignment accuracy in the augmented reference. We now propose multiple variant models, each aiming to give higher scores to variants that will impart a greater net benefit when considering accuracy and blowup. The window size *s* is used in three separate places in the software (described below) and should typically be set to the maximum read length.

#### 3.1.1 Variant models

Let *G*_*ref*_ denote the linear reference genome and *G\** the complete augmented genome including all variants in the population. Let *G* be a possible result of a VIS, i.e. an augmented genome that includes a subset of population variants. For simplicity, assume all variants are SNVs (substitutions). Let a *localized s-mer* ⟨*s, l*⟩ be a string of length *s* (the configurable windows size) that matches some combination of alleles in an augmented genome *G* starting at offset *l*; we also call these simply ⟨*s, l*⟩-mers. For instance, if *G* is `GATYACA`, where `Y` can be either `C` or `T`, then ⟨`GAT`, 0⟩, ⟨`TCA`, 2⟩ and ⟨`TTA`, 20⟩ are all ⟨3, *l*⟩-mers of *G*. For an ⟨*s, l*⟩-mer *σ*, let *p*(*σ*) be the probability a random ⟨*s, l*⟩-mer drawn from a random individual in the population equals *σ*. This can be calculated as:

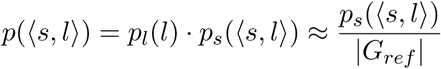

where *p*_*l*_(*σ*) is the probability a random *s*-mer begins at *σ*’s offset, which we approximate by 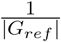. *ps*(*σ*) is the probability a localized *s*-mer starting at *l* has alleles matching *σ*’s. We approximate *p*_*s*_(*σ*) by assuming independence and multiplying the frequencies of each allele, or, if phasing information is available, by using allele co-occurrence frequencies.

##### Population Coverage

The *population coverage C*(*G*) of an augmented reference *G* is proportional to the population variation included, weighted by allele frequency. Specifically:

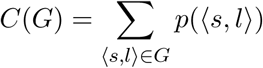

Note that *C*(*G*_*ref*_) ≤ *C*(*G*) ≤ *C*(*G\**) = 1.

We want to prioritize alleles according to how much they increase *C*(*G*). To do so accurately, each variant’s effect on *C*(*G*) must be calculated according to which nearby variants (within *s−* 1 positions) are already in *G*. While this is possible, it requires much recalculation of scores as variants are added to *G*. It also means there is no way to produce a single, static list of per-variant model scores. For these reasons, we instead compute each variant’s effect on *C*(*G*) assuming that all surrounding variants are already in *G*; in other words, we compute the decrease in *C*(*G*) caused by removing the variant from *G*\*. We call this the *complete graph* assumption.

Although FORGe is capable of using phasing data — describing which alleles co-occur on the same haplotype — the complete graph assumption makes this irrelevant for our calculation here. We do make (optional) use of phasing data in the *Hybrid* model, discussed below.

##### Uniqueness

The *uniqueness U* (*G*) of a genome *G* decreases as the multiplicities of its *k*-mers increase, i.e. as the genome becomes more repetitive. Let *f*_*G*_(*s*) be the number of ⟨*s*′, *l*′⟩-mers in genome *G* with *s* = *s*′. We define uniqueness of the genome as:

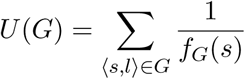

Adding a variant to the genome can either increase or decrease *U* (*G*). Specifically, an ⟨*s, l*⟩-mer overlapping the variant increases *U* (*G*) if there is no other ⟨*s*′, *l*^′^⟩-mers with *s* = *s*′. Alternately, an ⟨*s, l*⟩-mer overlapping the variant decreases *U* (*G*) if there are anyother ⟨*s*′, *l*^′^⟩-mers with *s* = *s*′.

While we rely on this definition below, we do not expect uniqueness alone to be an effective variant model. This is because for most variants all the added (overlapping) ⟨*s, l*⟩-mers are unique. All such variants therefore receive an identical score. The hybrid measure, presented next, effectively breaks ties by also considering allele frequency.

##### Hybrid score

The *hybrid score H*(*G*) of a genome *G* considers both population coverage and uniqueness. Again let *fG*(*s*) be the number of ⟨*s*′, *l*^′^;⟩-mers in *G* with *s* = *s*′ and let *p*⟨*s*′, *l*′⟩ be the probability a random ⟨*s, l*⟩-mer drawn from a random individual equals⟨*s*′, *l*′⟩. We define the hybrid measure *H*(*G*) of an augmented reference *G* as

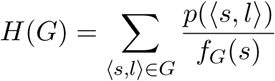

Note that this is simply the dot product of the terms from the *C*(*G*) and *U* (*G*) sums.

For a variant *v*, we wish to compute the increase in *H*(*G*) caused by adding *v*. For each *s, l*⟩-mer overlapping *v* and containing the alternate allele, let ⟨*s, l*_1_⟩, ⟨*s, l*_2_⟩, *…,* ⟨*s, l*_*n*_⟩be all other ⟨*s, l*⟩-mers with the same sequence *s*. Before adding *v*, the hybrid score can be written as

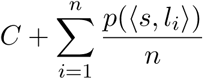

where *C* is the hybrid-score portion due to the ⟨*s*^′^;, *l*⟩-mers with ⟨*s*⟩ ^′^ ≠ *s*. After adding ⟨*s, l*⟩ to *G*, the score becomes

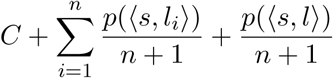

The change in hybrid score due to the addition of ⟨*s, l*⟩ is

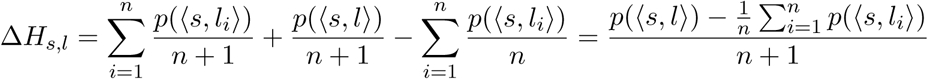

Assuming each ⟨*s, l*⟩-mer overlapping variant *v* has a distinct sequence *s*, their Δ*H*_*s,l*_ terms are independent. Thus the total change in hybrid score due to the addition of *v* is the sum of the Δ*H*_*s,l*_’s for each ⟨*s, l*⟩-mer overlapping and including *v*.

There are a couple caveats to how FORGe implements the hybrid model. First, As with the *Pop Cov* model, we make the complete graph assumption, allowing us to produce a scored variant list without dynamic re-scoring of variants as they are added. Second, computing Δ*H*_*s,l*_’s for all variants is expensive, since it involves calculating the read probability for each other occurrence of sequence *s* for every overlapping ⟨*s, l*⟩-mer. Instead, we approximate it using average probabilities. Specifically, we pre-calculate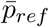, the average *p*(⟨*s, l*⟩) for all ⟨*s, l*⟩-mers in *G*_*ref*_, and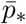, the average *p*(⟨*s, l*⟩) for all ⟨*s, l*⟩-mers in *G\** but not in *G*_*ref*_. We approximate the summation with a weighted average:

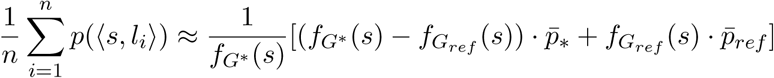

Whereas the complete graph assumption rendered phasing data irrelevant to the *Pop Cov* model, we can use phasing data in the *Hybrid* model. This is because the *Hybrid* model weights the terms of the sum according to their frequency in the genome. By default, FORGe uses phasing information when it is available.

##### Hybrid score implementation

The Uniqueness and Hybrid models are concerned with *s*-mer counts both in the linear reference genome (*G*_*ref*_.) and in the complete augmented reference (*G\**). FORGe uses Jellyfish v2.2.6 [42] to calculate these counts. Since Jellyfish counts *s*-mers in a FASTA input file, FORGe must first construct an augmented FASTA such that ⟨*s, l*⟩-mers in this FASTA map one-to-one to ⟨*s, l*⟩-mers in *G\**. This is also the goal of the Enhanced Reference Genome [7] representation, which accomplishes this by adding 2^*k*^ *−*1 “enhanced segments” for every length-*s* window containing *k* variants. Thus, to obtain *s*-mer counts for *G\**, we first constructed such as FASTA file using our implementation of the ERG, then counted *s*-mers using Jellyfish.

Once *s*-mers have been counted, FORGe computes the average probability for reads in the linear reference 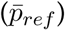 and in the complete augmented reference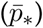, for use in the *Hybrid* model formula. Finally, we compute the change in *H*(*G*) for each *s*-mer in both and update the *Hybrid* model scores for every variant with an alternate allele in that read. After this, we have the full set of *Hybrid* model scores for all variants.

##### Considering blowup

Adding variants to the augmented reference increases computational costs, including (a) size of the index on disk, (b) memory footprint during read alignment, and (c) time required for read alignment. We collectively refer to these as “blowup.” Blowup is most drastic in genomic regions where variants are densely clustered, driving a exponential increase in the number of allelic combinations possible. A model based purely on minimizing blowup would prioritize isolated variants over those in clusters. We do not expect such a model to perform well on its own, though, since (like the *Uniqueness* model described above) it would fail to prioritize among the isolated variants. For this reason, we sought a way to combine a blowup avoidance strategy with the models already described above.

##### Selecting variants with blowup avoidance

After ranking variants, FORGe selects the subset of variants to include in the augmented reference. The user specifies either a number or a fraction of all variants to include. In the simplest case, variants are chosen in order, starting with the highest-scoring variant, until the desired number have been included.

As an additional defense against blowup, we also propose a dynamic re-scoring scheme that can be added to an existing model. In this scheme, when a variant is added to the reference, FORGe searches for other variants within *s* nt (the window length) of the added variant that have not yet been selected for addition. These nearby variants are re-scored by multiplying their score by a penalty factor *w*, where 0 < *w ≤*1. By letting *w* be variable, FORGe can trade off between maximizing the model score and minimizing blowup. *w* = 1 maintains the original scores, whereas a penalty near *w* = 0 would ensure all isolated variants were added before any neighboring variants. We found that a penalty of *w* = 0.5 performed well in practice, and this is FORGe’s default, used in all experiments performed here. *Pop Cov+* and *Hybrid+* are how we refer to those models when they are combined with this dynamic re-scoring scheme.

##### Breaking ties

Variants can be given identical scores using these models. For instance, variants with the same allele frequency will receive the same score by the *Pop Cov* model. These ties are broken according to the variants’ position on the genome. We define variant A to be upstream of (and higher priority than) variant B if it is on a lower-numbered chromosome or if its offset is to the left of B’s.

## 4 Discussion

FORGe’s modeling of positive and negative effects of including genetic variants in an augmented reference yields accuracy-blowup tradeoffs superior to current approaches. We proposed models for prioritizing variants with distinct rationales and strengths. We found repeatedly that the most advantageous set of variants consisted of a fraction (6– 30%) of the variants called in the 1000 Genomes project. This was true across a variety of alignment scenarios: for two different graph alignment methods (HISAT2 and ERG), for both unpaired and paired-end alignment modes, when just SNVs and when both SNVs and indels are included, for both a single human chromosome and for the whole human genome, and for both CEPH and YRI individuals. We also showed that FORGe’s modeling can substantially improve downstream results related to reference bias and HLA typing, also at relatively low variants-inclusion fractions.

To test if FORGe’s results yield a simple filtering rule, we can translate the peak-performing variant inclusion fractions for the *Pop Cov* model into allele frequency thresholds. For the Chromosome-9 NA12878 experiments, the 8% variant inclusion fraction performed best in both the unpaired (Figure 1) and paired-end (Supplementary Figure 3) experiments. This translates to an allele frequency threshold of ≥7.42%. For the Chromosome-9 YRI experiment (Supplementary Figure 4), the best-performing fraction of 10% translated to an allele frequency threshold of ≥ 3.76%. While these fall on either side of the 5% threshold used in at least one prior study [13], further work is needed to establish whether that or any other simple threshold is justifiable in general. A finer-grained sweep over variant inclusion fractions would yield a sharper threshold, for example. For now, our strategy of gradually introducing variants in the context of a simulation study is both principled and practical.

FORGe and HISAT2 combine to make a practical graph aligner that works with human data with large variant databases like the 1000 Genomes Phase 3 call set. Using `hisat2-build` to index a GRCh37-based graph genome with the top 8% of variants from Phase-3 set required 4 hours and 165 GB of memory. Aligning 20 million reads to this graph required 19 minutes and 6.5 GB of memory, about 50% more time and 50% more memory than HISAT2 requires to align to the linear GRCh37 genome. (To prioritize the variants prior to indexing, FORGe required about 110 minutes on a single processor.) This is competitive with the performance of aligners like Bowtie 2 and BWA-MEM when aligning to the linear reference, suggesting graph-based tools are ready for broader use.

Though we estimate that the overall improvement in alignment accuracy for a 40x whole-genome DNA sequencing experiment would lead to 4.8M more correctly aligned reads and 1.2M fewer incorrectly aligned reads, the magnitude of the improvement imparted by modeling variants depends on the genomic region. For some regions and variant classes (rare, isolated SNVs), the benefit is small. To improve alignment to these regions might require an iterative approach that aligns to a graph containing known variants, calls donor-specific variants, then realigns to a graph that includes both. Strategies like this are implemented in the GATK HaplotypeCaller [20], GraphTyper [21] and other tools [43]. Better variant models might also benefit these hard cases. Even so, the effects we measured translate into substantial net increases in the number of correctly aligned reads, and the results are pronounced in regions such as MHC as shown by Supplementary Figures 6 and 9.

An ethnicity-specific reference conferred a slight accuracy improvement compared to a pan-ethnic reference with a similar number of variants. This is notable in light of proposals to use ethnicity-specific references [36, 37]. It suggests that the advantages of an inclusive reference, applicable regardless of the donor individual’s ethnicity, might outweigh the slight accuracy gain that comes with ethnicity-specificity. Also, ethnicity-specific references could be counterproductive or misleading in cases where donor ethnicity is reported incorrectly or where the donor is admixed [44].

The accuracy achieved at relatively small fractions of the 1000 Genomes variants has implications for the design of graph aligners. A central challenge for these tools is to operate efficiently even when variants are densely clustered, causing local explosion in the number of allelic combinations. But our observations that peak accuracy occurs at a relatively small fraction of variants, and that memory footprint increases by a factor of 2 or less at peak accuracy, suggests that this is not a major barrier to practical graph-genome alignment as long as variants are chosen carefully.

It should also be possible to adapt FORGe to study how including structural variants can improve alignment. A common observation of studies that have assembled human genomes from long reads is that the assemblies contain many megabases of sequence not present in the standard human reference [37, 38, 45, 46]. The models we propose are equally applicable to structural variants, assuming the variants are called in enough individuals to estimate allele frequencies accurately.

While we primarily investigated unpaired alignment here, we also showed that the chromosome-9 results generalized to paired-end alignment (Supplementary Figure 3). Generally speaking, this work can be adapted to paired-end alignment, with the main issue being how to adjust the method’s window lengths as a function of the paired-end dataset’s read and fragment lengths. The windows in question are (a) the *s*-mer length used in the model, (b) the maximum window length used when forming enhanced segments for ERG-based alignment, and (c) for blowup avoidance in the *Pop Cov+* and *Hybrid+* models, the radius to look within when seeking nearby variants to deprioritize. While one option is to simply increase the window size to the maximum fragment length, that can easily lead to an unacceptable blowup penalty. For this reason, we suspect that there is no practical way to adapt our ERG-based to paired-end alignment; rather, as we explain, we think it is best viewed as a model for the seed-finding step of a seed-and-extend aligner that might itself handle paired ends. Initial experiments suggest that it is practical to leave the window lengths relatively short — the length of a read rather than a fragment — when using HISAT2 for paired-end alignment (Supplementary Figure 3). Further exploration is needed to more fully characterize the relationship between FORGe window length and fragment and read lengths for paired-end reads.

## 5 Acknowledgments

We thank Daehwan Kim and Michael Schatz for their helpful comments on a draft of this manuscript.

## 7 Availability of data and materials

The FORGe software is available at: https://github.com/langmead-lab/FORGe/releases/tag/v1.1 under the open source MIT license. The FORGe repo also includes our implementation of the Enhanced Reference Genome, originally proposed by Satya et al [7]. The experiments described in this paper are open source under the open source MIT license and available at https://github.com/langmead-lab/FORGe-experiments. We provide three variant datasets and HISAT2 indexes, available at ftp://ftp.ccb.jhu.edu/pub/langmead/forge. These include the optimal graph genomes described above for chromosome 9 using only SNVs (`NA12878_chr9` subdirectory), for chromosome 9 using SNVs and indels (`NA12878_chr9_indels` subdirectory), and for the full genome (`NA12878_full_genome` subdirectory), all with NA12878 and family members withheld. We also provide what we consider the best full genome graph, derived from all1000 Genomes SNVs and indels from all individuals and all populations (`best_full_genome` subdirectory). The variants and index files in the (`best_full_genome` subdirectory) are the ones we suggest others use for future studies.

## 8 Authors’ contributions

JP and BL designed the method and wrote the software and manuscript.

## 9 Ethics approval and consent to participate

No ethics approval was required for this work.

